# The construction and operation of type IV pili impose a variable energetic burden across phylogenetically distant bacteria

**DOI:** 10.64898/2026.09.14.751554

**Authors:** Ahmed O. Yusuf, Zil K. Modi, Matthias D. Koch

**Author notes:** Equal contribution (Ahmed O. Yusuf, Zil K. Modi).

## Abstract

Type IV pili (T4P) are dynamic surface appendages that mediate essential biological functions and virulence traits, yet their energetic burden on cellular budgets in light of fluctuating host environments remains unexplored. Here, we present a comprehensive economic analysis of T4P construction and operation in ATP equivalents, following established frameworks of flagella analyses.

Using *Pseudomonas aeruginosa* as a model system, we quantify the total cellular burden to synthesize the T4P machinery, maintain the inner-membrane pool of major pilin (PilA), and drive repeated cycles of pilus extension and retraction over a generation. We estimate that the T4P system consumes ~0.7% of the total cellular energy budget, dominated by PilA monomer production. Conversely, the operational cost of dynamic T4P fibers is negligible due to their intermittent activity - contrasting sharply with the high continuous cost of rotating a polar flagellum.

Extending this framework across five phylogenetically diverse species (*P. aeruginosa, Vibrio cholerae, Caulobacter crescentus, Neisseria* spp., and *Myxococcus xanthus*) reveals that T4P investment varies tenfold (0.2–1.8% of the cellular budget), driven by differences in pilin size, machine number, pilus extension rates, and cell volume. *Neisseria* is a distinct outlier whose high extension rate makes operational costs approach construction costs, while in all other species construction dominates. These findings indicate that changes in nutrient availability or surface association may modulate pilus number and length as a strategy to optimize energetic burdens during host-pathogen interaction.

## Introduction

Type IV pili (T4P) are dynamic, retractable filamentous appendages expressed by a wide range of Gram-negative and some Gram-positive bacteria. T4P mediate many biological functions, ranging from surface interaction (including adhesion, motility, micro colony formation, and sensing) to phage infection and natural transformation [1–8]. Their expression and dynamics are tightly regulated by surface contact, nutrient availability, cyclic-di-GMP levels, and mechanical cues [9–13].

In contrast to the well-quantified energetic costs of flagellar systems, the bioenergetic investment required for T4P assembly and operation has remained largely unexplored. Construction of a single flagellum typically requires ~10^7^-10^8^ ATP, while continuous highspeed rotation (100–300 Hz) driven by a large proton flux (~1,200 protons per revolution) can consume tens of thousands of ATP equivalents per second per motor [14]. Across the tree of life, flagella impose substantial and highly variable costs across species, with energetic investment ranging from <0.1% to >40% of the total cellular energy budget depending on cell size, flagellar number, and motility regime [14].

*Pseudomonas aeruginosa* typically produces one polar flagellum together with a variable number of T4P. Different strains and environmental conditions lead to markedly different piliation levels. Laboratory strains often display modest piliation in liquid culture, whereas surface-attached cells, biofilms, or retraction-deficient (Δ*pilT*) mutants can assemble dozens of machines [8, 15]. Clinical isolates frequently exhibit altered T4P dynamics that correlate with virulence and persistence [16–18].

Across other species, T4P dynamics vary considerably. *Myxococcus xanthus* for example assembles 5–10 T4P at the leading cell pole during S-motility, similar to *Pseudomonas* [19–21]. In contrast, *Neisseria* species typically display 5–7 pili per cell on average, but production rates reach ~196 pili per minute in *N. gonorrhoeae* [22–25]. Other species like *Vibrio cholerae* employ several distinct T4P systems (for example MSHA and competence pili) with highly dynamic extension and retraction [26–28], while *Caulobacter crescentus* and *Acinetobacter baylyi* produce fewer and more tightly regulated pili [29, 30].

To determine whether these differences reflect energetic constraints, we performed a detailed bioenergetic accounting of T4P construction (machine proteins and major pilin pool) and operation (extension/retraction powered by PilB and PilT) in *P. aeruginosa* using cysteine-maleimide labeling data [31]. We then extended the same quantitative framework across phylogenetically diverse species to test whether relative T4P investment is evolutionarily constrained and how nutrient conditions may drive observed strain- and species-specific differences in pilus number and length.

## Results

To estimate the bioenergetic costs associated with type IV pili (T4P), we separately calculated the construction costs of the pilus machine, the intracellular PilA monomer pool, and the operating costs of pilus extension and retraction. All costs are expressed in ATP equivalents, following the framework of Schavemaker and Lynch used to perform a similar calculation for the energy budget of flagella [12]. We used a conversion factor of 29 ATP per amino acid, which accounts for both the direct cost of peptide bond formation and the opportunity costs of amino acid biosynthesis [32]. Protein amino acid (AA) lengths were obtained from the pseudomonas.com database [33].

### Construction cost of one T4P machine

In *Pseudomonas*, the core structural T4P machine complex consists of the PilMNOPQ complex, the alignment protein PilF, the inner membrane platform PilC, and several minor pilins that are thought to form one priming complex inside each functional T4P machine [1, 2, 15]. Many of these proteins form multimeric structures in their functional state [34]. Further, all pilins are processed by the peptidase PilD. Since we are unaware of a good estimate for the copy number of PilD per cell, we included one copy per machine for the sake of completeness.

As shown in Table 1, the total cost to make one T4P machine complex is roughly 613,000 ATP. The largest cost of assembling the core machine complex is the formation of the outer membrane pore PilQ, requiring nearly half of the total construction cost of one machine. The total number of machines in individual *P. aeruginosa* cells is not well known. To understand how many functional T4P machines there are in one cell, we fluorescently labeled pilus fibers in a retraction deficient *pilTU* deletion mutant (see Supplementary Results). Here, functional machines can be identified because they extend a pilus once but the pilus cannot be retracted, hence, every functional T4P machine is piliated. We observed a broad distribution of 0 – 9 pili per cell, with a mean of 3 pili per cell (Figure 1A). However, cryo-electron microscopy data of surface grown cells have revealed that individual cells can have as many as 80 T4P machines on one pole (albeit only a small fraction appeared functional) [15].

**Table 1.**
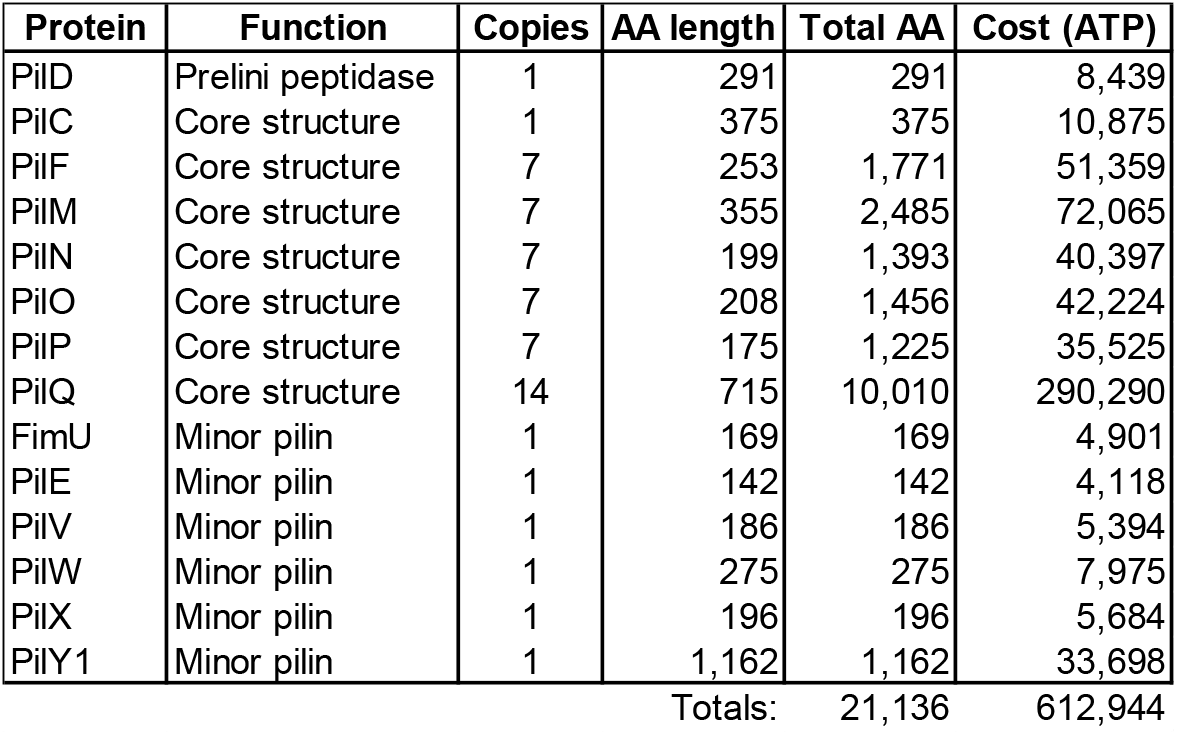
Cost to generate the individual components necessary to make one functional TFP machine.

**Figure 1.**
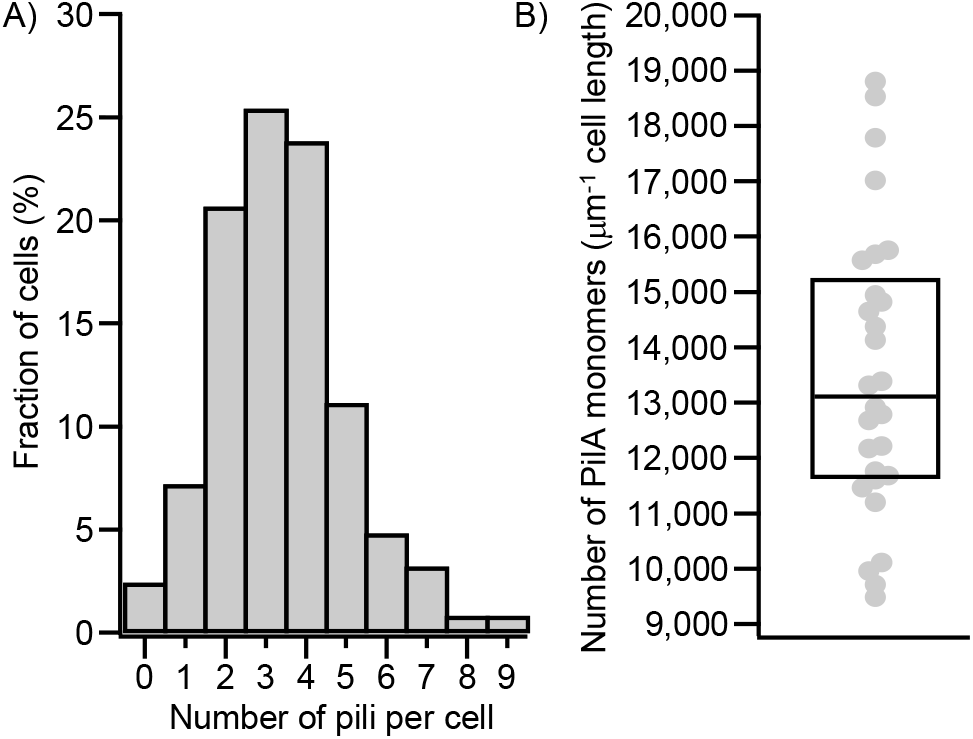
Estimate of number of TFP machines (A) and number of PilA monomers (B) per cell. A) The number of machines was estimated by counting the number of pili in a non-retracting *pilTU* motor mutant. **B)** The number of PilA monomers was estimated based on changes in the total cell body fluorescence during extension of pili.

### Construction cost to make the extension and retraction ATPases PilB and PilT

Extension and retraction of pilus fibers is mediated by the hexameric ATPases PilB and PilT. Although their exact copy number in a single cell is not known, it is reasonable to assume that at least one hexameric copy of each ATPase is required for one functional T4P machine, equating to roughly 159,000 ATP. In reality, there are likely many more copies in a cell, presumably at least a 10-fold excess. This shows that the energetic burden to make enough of the ATPases required to operate one machine is likely in a similar range as assembling the machine complex itself.

**Table 2.** Cost to generate one extension and one retraction motor complex (hexamer) necessary to make one functional TFP machine.

| Protein | Function | Copies | AA length | Total AA | Cost (ATP) |
| --- | --- | --- | --- | --- | --- |
| PilB | ATPase | 6 | 567 | 3,402 | 98,658 |
| PilT | ATPase | 6 | 345 | 2,070 | 60,030 |
| Totals: |  |  |  | 5,472 | 158,688 |

### Generating the inner membrane pool of the major pilin PilA

PilA is a small protein with 143 AA in *P. aeruginosa*, equating to roughly 4100 ATP needed to generate one monomer. The density of PilA in a pilus fiber is roughly 1 PilA/nm, requiring several hundreds to thousands of monomers to build a typical filament [34, 35].

However, the incorporation of PilA into the fiber relies on diffusion of these molecules in the inner membrane to the T4P machine complex [8]. For this process to happen efficiently, meaning monomers are readily available for the ATPase to incorporate at the maximum rate of ATP turnover, the concentration of PilA in the inner membrane needs to be much higher than the number of molecules that end up going into a fiber. To estimate the concentration of PilA monomers in the cell envelope, we used fluorescent staining of the monomer and measured the reduction of brightness in the cell envelope relative to the length of an assembled/extended pilus (see Supplementary Results and Methods). Based on this analysis, we estimate that a typical cell has a density of approximately 13,000 PilA monomers per micrometer of cell length (Figure 1B). The average cell length is about 3 μm, equating to roughly 39,000 PilA monomers per cell. This brings the total estimated construction cost of the PilA pool to ~162 million ATP per cell, representing the major construction cost of the T4P system, exceeding the cost of the machines and motors by at least one order of magnitude.

**Table 3.**
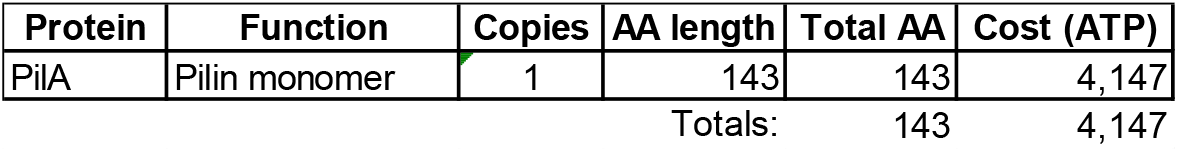
Cost to generate one pilin monomer. Note, many hundred to thousand are polymerized to form a fiber and many tens of thousands are required in the inner membrane to facilitate the extension mechanism.

| Protein | Function | Copies | AA length | Total AA | Cost (ATP) |
| --- | --- | --- | --- | --- | --- |
| PilA | Pilin monomer | 1 | 143 | 143 | 4,147 |
| Totals: |  |  |  | 143 | 4,147 |

### Comparison of construction cost components

As explained above, the absolute number of machine complexes, ATPases, or pilin monomers is not well characterized, and we have attempted to give reasonable estimates for *P. aeruginosa*. To allow for a fair future comparison, especially due to changes in T4P expression between different conditions such as nutrient environments, mutant backgrounds, etc, we modeled the total ATP required for each of the three components as a function of their total copy number (Figure 2). This analysis shows that in realistic cases (moderate amount of T4P machines and motors), the major contribution to the energy requirement is the generation of pilin, followed by the generation of T4P machines, and lastly the generation of the ATPases. Only in cases with extremely high T4P machine count (such as in surface grown cells) and low PilA monomer pool these contributions might approach each other on an energetic basis.

**Figure 2.**
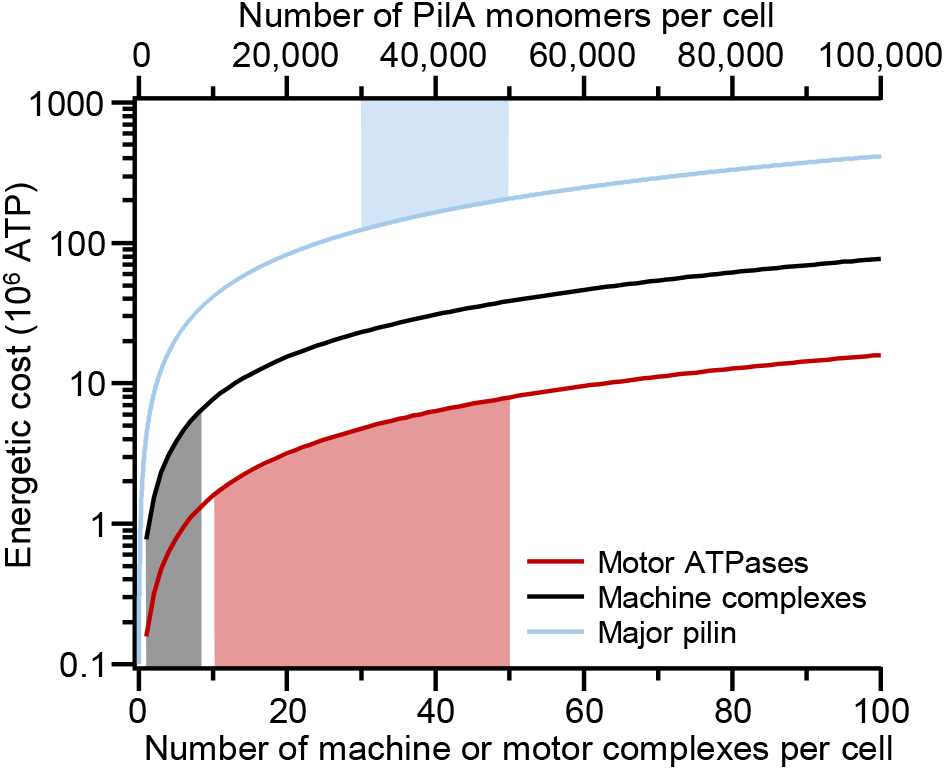
Comparison of the total construction costs of T4P machines, motor complexes, and the pilin pool during the life cycle of a cell of 30 minutes. Shaded areas indicate estimated ranges for typical *P. aeruginosa* cells grown under standard liquid laboratory conditions (LB medium, mid-log, 37C shaking).

### Operating (maintenance) cost of pilus extension and retraction

Cysteine-maleimide labeling experiments of T4P fibers has revealed that the number and length of pili produced by individual cells follow a exponential distributions with a mean of 8 pili per cell per minute a mean length of 0.8 µm [36]. Over a typical 30-minute cell growth cycle, the average total pilus fiber length extended per cell is therefore 8 × 30 × 0.8 µm = 192 µm.

Extension and retraction by PilB and PilT require ATP. Here, we assumed 2 ATP per PilA subunit incorporated or removed [37, 38], equivalent to 4,000 ATP per micrometer of pilus for one complete extension-retraction cycle of a 1 µm long pilus. Consequently, the average operating cost of T4P dynamics over one complete cell cycle is approximately 768,000 ATP per cell.

Because both the number of pili and their lengths are exponentially distributed, the total operating cost across a population of cells is also exponentially distributed. This results in substantial cell-to-cell variation (Figure 3). The 10th percentile of least active cells expends approximately 110,000 ATP on pilus extension and retraction during one cell cycle, while the 90th percentile of most active cells expends approximately 1,650,000 ATP. This ~15-fold difference between the lowest and highest deciles highlights that a small subpopulation of cells incurs a disproportionately high energetic burden for pilus dynamics. However, compared to the total construction cost of the T4P system, operating T4P fibers is about two orders of magnitude cheaper than generating the individual components.

**Figure 3.**
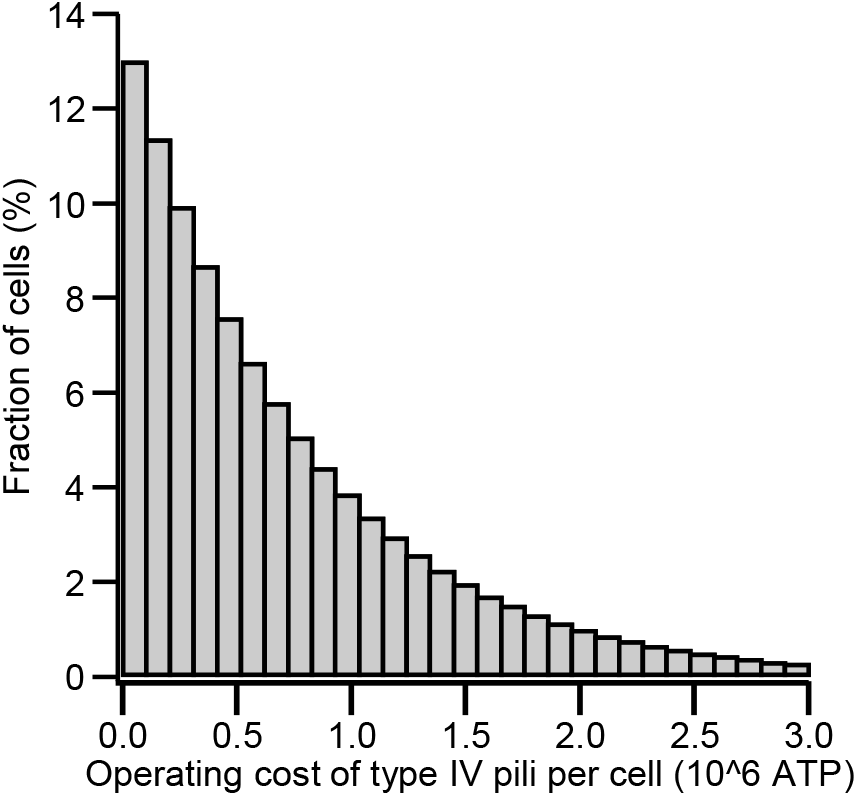
Operating cost to extend pili for a population of cells during one full replication cycle of a cell of 30 minutes.

### Comparison to the total cellular energy budget

To place the cost to construct and operate T4P into the broader cellular energy budget, we compared them to the total cost of building and operating one complete *P. aeruginosa* cell over one division cycle. Using the bioenergetic framework of Lynch and Marinov [32], the lifetime energetic requirement of a cell is the sum of the biosynthetic cost of cell growth and a basal maintenance cost that accumulates over the generation time. Both costs scale nearly linearly with cell volume. Treating an average *P. aeruginosa* cell as a 3 × 0.6 µm rod gives a volume of roughly 0.85 µm^3^. Relative to the reference bacterium in Lynch and Marinov’s framework (*E. coli* with ~1 µm^3^ volume that requires ~1.6 × 10^10^ ATP), a 30-minute generation time for *Pseudomonas* yields a total cellular energy budget of approximately 2.4 × 10^10^ ATP per generation.

This shows that the T4P system accounts for only ~0.68% of the cell’s total energy expenditure (Table 4). The only significant expense for the cell is the creation of the major pilin pool, while all other T4P costs are negligible.

**Table 4.** Comparison of T4P costs to total cellular budget. This table assumes three T4P machines and 30 ATPase hexamers each (10x excess).

|  | Cost (ATP) | Cost (%) |
| --- | --- | --- |
| Machine construction | 1,838,832 | 0.008% |
| ATPase constuction | 4,760,640 | 0.020% |
| Pilin pool | 161,733,000 | 0.674% |
| Operating cost | 768,000 | 0.003% |

### Comparison to the cost of flagella

To put the energetic cost of the T4P system into perspective, we compared it with the cost of producing and operating the single polar flagellum of *P. aeruginosa*, using the detailed estimates provided by Schavemaker and Lynch (Table 5) [14]. The construction of one flagellum requires approximately 232 million ATP, which exceeds the average total cost of the entire T4P system of 168 million ATP. When the operating costs of flagellar rotation are included, the total investment in the flagellum is estimated to be three-to four-fold higher than that of the T4P system, and averages ~10% of the total energy budget across different bacterial species [14].

**Table 5.**
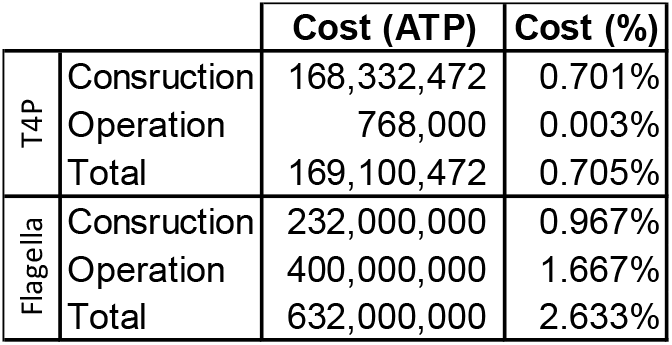
Comparison of cost of T4P vs flagella compared to the total cellular energy budget.

|  |  | Cost (ATP) | Cost (%) |
| --- | --- | --- | --- |
| T4P | Construction | 168,332,472 | 0.701% |
|  | Operation | 768,000 | 0.003% |
|  | Total | 169,100,472 | 0.705% |
| Flagella | Construction | 232,000,000 | 0.967% |
|  | Operation | 400,000,000 | 1.667% |
|  | Total | 632,000,000 | 2.633% |

These results show that, on average, *P. aeruginosa* invests significantly more energy into its flagellum than into its T4P. The much higher operating cost of the flagellum compared with T4P stems from fundamentally different modes of action. The flagellar motor rotates continuously at high speed (typically 100–300 Hz in *P. aeruginosa* under load [39] and is powered by a large proton flux of approximately 1,200 protons per revolution (or roughly 1.2–3.6×10^5^ protons per second per motor) [40]. Assuming ~4 protons per ATP synthesized by ATP synthase under typical cellular conditions, this proton flux translates into an operating cost on the order of ~6.6×10^4^ ATP per second per flagellum [14]. Over a 30-minute generation time, flagellar rotation therefore consumes several hundred million ATP.

In contrast, T4P operate in short, intermittent extension–retraction cycles with a low duty cycle. Even with the experimentally observed exponential distribution of pilus production, the average operating cost remains only 768,000 ATP per generation — roughly two orders of magnitude lower than continuous flagellar rotation. Thus, while the PilA monomer pool makes the construction cost of T4P comparable to that of one flagellum, the *operating* cost of pili is far smaller because they function episodically rather than continuously propelling the cell through liquid.

### Strain specific variation

Across phylogenetically diverse bacteria, the number of T4P machines per cell, pilus production rates, and average pilus lengths vary modestly, with a few notable exceptions.

Most species assemble approximately 2–5 active machines per cell and maintain a small number of pili (typically 1–10 per minute) with average lengths of 0.8–1.5 µm [8, 19, 20, 23, 27, 28, 41, 42]. Notable outliers are *Pseudomonas aeruginosa* which can produce dozens of machines in certain surface-attached conditions or retraction-deficient mutants [15, 43] and *Neisseria* species that exhibit exceptionally high pilus production rates (~196 pili per minute) [26].

To compare T4P related energetic costs between species, we 1) assumed a conserved major pilin monomer density per unit area in the inner membrane (scaled from our direct cysteine-maleimide measurements), 2) included the same estimate of T4P construction cost per machine from above and 3) assumed a similar packing density of pilin monomers in the fiber. We further included typical growth rates under laboratory conditions [44–48], the differences in pilin size (45–208 amino acids) [49–52], cell geometry and size [8, 53–57], and T4P dynamics in the analysis [8, 19, 20, 23, 27, 28, 41, 42] (Table 6). Based on this analysis, we found that the total construction cost varies one order of magnitude between 40 – 350 million ATP while the operating cost varies more than two orders of magnitude between 0.16 – 59 million ATP during the life cycle of a cell (Table 7). Similarly, the total energetic burden relative to the cellular energy budget varies ~7.5-fold between 0.23 – 1.8%. However, the relative energy expenditure on T4P construction vs operation differs markedly by species: while *V. cholera* spends only 0.1% of the T4P related energy cost on operation of filaments, *Neisseria spp*. spend almost equal amounts on both due to extremely high pilus production rates.

**Table 6.**
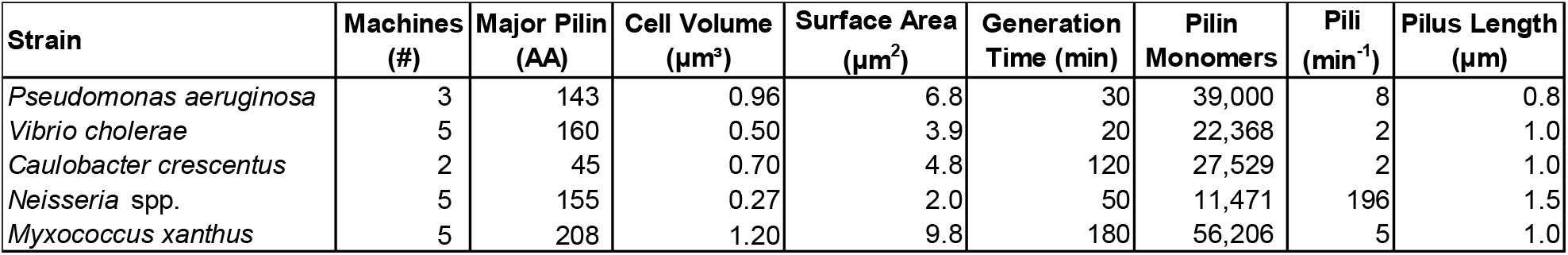
Input parameters required to model relative energetic costs of constructing and operating T4P between different species.

**Table 7.** Comparison of total T4P cost and relative contribution to the total cell budget between common T4P-expressing strains.

| Strain | Construction Cost (ATP) | Operating Cost (ATP) | Total Cost (%) | Operating to Construction Cost (%) |
| --- | --- | --- | --- | --- |
| *Pseudomonas aeruginosa* | 168,342,000 | 768,000 | 0.705 | 0.5 |
| *Vibrio cholerae* | 114,800,882 | 160,000 | 0.920 | 0.1 |
| *Caulobacter crescentus* | 40,331,882 | 960,000 | 0.236 | 2.4 |
| *Neisseria* spp. | 62,575,294 | 58,800,000 | 1.798 | 94.0 |
| *Myxococcus xanthus* | 350,048,882 | 3,600,000 | 1.179 | 1.0 |

## Discussion

Our analysis shows that the energetic investment in T4P is relatively small, accounting for ~0.7% of the cellular energy budget in *P. aeruginosa* and remaining consistently low (~0.2-1.8%) among several phylogenetically diverse species. This contrasts sharply with flagellar systems, which often require >10% and reach >40% of the cellular budget to sustain continuous high-speed rotation [14]. This economy stems primarily from the low operating costs of T4P: their intermittent, low-duty-cycle dynamics keep ATP consumption for extension/retraction modest compared with the relentless proton flux of flagellar motors.

Interestingly, the efficiency of motility (costs per distance traveled) is similar in magnitude (~10^3^ ATP per µm) for both systems, based on an instantaneous twitching speed of ~0.3 µm/s for retraction of an individual pilus and continuous swimming at ~60 µm/s [58, 59]. This similarity is striking given that the two modes of motility operate under very different boundary conditions: T4P must overcome intermittent surface friction and adhesion while flagella work against continuous viscous drag. However, many T4P retract without attaching and therefore do not contribute to translocation [60], so the time-averaged efficiency of twitching over minutes to hours is likely lower than that of continuous flagellar swimming.

**Table 8.** Efficiency of flagella vs T4P mediated motility.

|  | Speed<br>(μm/s) | Consumption<br>(ATP/s) | Efficiency<br>(ATP/μm) |
| --- | --- | --- | --- |
| T4P | 0.3 | 427 | 1,422 |
| Flagellum | 60 | 66,400 | 1,107 |

What are the energy-economic implications for T4P regulation? Recent findings suggest that in *P. aeruginosa*, both T4P count and filament length are regulated transcriptionally by modulating the expression of all core T4P genes via cAMP [8, 13, 61, 62]. This is consistent with surface-attached cells and biofilm-grown cells exhibiting increased piliation and altered machine distribution compared to planktonic cells [63, 64]. Surface contact has also shown to trigger engagement of the T4P motors to initiate retraction, although this behavior has not been consistently observed by all studies [41, 60, 65]. Similarly to surface grown cells of *P. aeruginosa, Thermus thermophilus* maintains a large excess of assembled machines (~33 per cell) relative to actively piliated ones (~6 per cell), suggesting a potential reservoir whose activity could be regulated based on environmental cues [15, 66]. Rather than controlling the generation of new machines, both *Myxococcus xanthus* and *P. aeruginosa* dynamically reposition the extension and retraction motors on pre-positioned bipolar machines using chemosensory systems [61, 67].

Consistent with our findings that the pilin pool presents the single largest energetic cost of the T4P system, modulation of PilA expression appears more widespread and dynamically utilized across species [68–70]. A recent study demonstrated that *Vibrio cholerae* modulates pilus dynamics by tuning the minor-to-major pilin stoichiometry [71]. Complementary work in *P. aeruginosa* established PilA abundance as a central lever for extension velocity and pilus length, and virulence traits including twitching motility, surface sensing, biofilm formation, and phage infection [72].

T4P energetics also interface directly with cellular energy status. Pre-assembled machines persist in stationary phase, yet low adenylate energy charge throttles motor activity, producing sparse, short pili [12]. Nutrients upshift rapidly reactivates dynamics without de novo synthesis, enabling biofilm dispersal while increasing susceptibility to pilus-dependent phages [12]. This “idling” strategy further reinforces the evolutionary advantage of maintaining a tunable PilA reservoir.

Overall, the constrained energetic footprint of T4P across bacteria suggests selection for efficient surface appendages that support multiple functions such as adhesion, motility, sensing, and gene transfer in variable environments without the high energetic burden imposed by other motility systems. Future studies should quantify how nutrient availability and surface association dynamically tune pilin pools, machine number, and T4P dynamics *in vivo*, and how these impact the energy economics of T4P.

## Acknowledgements

We would like to thank the entire Koch lab for stimulating discussion and the Department of Biology at Texas A&M for its supportive environment.

## Funding Statement

This work was supported by grant R35GM155280 from the National Institute of Health and startup funds from the College of Arts and Sciences, Division of Research, and Department of Biology at Texas A&M University to M.D.K, by the Nikon Center of Excellence and the Microscopy and Imaging Center (RRID:SCR_022128) at Texas A&M University.

## Author contributions

A.O.Y., Z.K.M, and M.D.K. designed research and wrote the manuscript. M.D.K. performed experiments and analyzed data.

## Competing Interest Statement

The authors declare no competing finical interests.

## Data Availability

Fluorescence microcopy movies of pilus dynamics are ~50 GB total and available from the authors upon reasonable request. All other data needed to evaluate the conclusions in the paper are present in the manuscript and/or the Supplementary Materials.

## Material and Methods

*Pseudomonas aeruginosa* PAO1 strain *pilA*-A86C *pilTU* [8] was grown overnight, backdiluted 1:500 and subsequently grown to mid log phase (OD ~ 0.1) at 37C with shaking in LB. Pilus fibers were fluorescently labeled as described previously [36]. In brief, mid log cells were incubated for 45 min with Alexa488-maleimide dye (Fisher), washed twice and then resuspended in 20 ul of fresh medium. 1ul of labeled cells were placed on a 1% agarose pad for imaging using a Nikon Ti2 microscope and a 100x NA1.45 Ph3 objective lens at 488 nm. Cell size, pilus length, and pilus fiber count were determined manually in ImageJ.

## Notes

### Competing Interest Statement

The authors have declared no competing interest.

## References

1. Craig, L., M.E. Pique, and J.A. Tainer, Type IV pilus structure and bacterial pathogenicity. Nat Rev Microbiol, 2004. 2(5): p. 363–78.

2. Burrows, L.L., Pseudomonas aeruginosa twitching motility: type IV pili in action. Annu Rev Microbiol, 2012. 66: p. 493–520.

3. Leighton, T.L., et al., Biogenesis of Pseudomonas aeruginosa type IV pili and regulation of their function. Environ Microbiol, 2015. 17(11): p. 4148–63.

4. Ellison, C.K., G.B. Whitfield, and Y.V. Brun, Type IV Pili: dynamic bacterial nanomachines. FEMS Microbiol Rev, 2022. 46(2).

5. Mattick, J.S., Type IV pili and twitching motility. Annu Rev Microbiol, 2002. 56: p. 289–314.

6. Gibiansky, M.L., et al., Bacteria use type IV pili to walk upright and detach from surfaces. Science, 2010. 330(6001): p. 197.

7. O’Toole, G.A. and R. Kolter, Flagellar and twitching motility are necessary for Pseudomonas aeruginosa biofilm development. Mol Microbiol, 1998. 30(2): p. 295–304.

8. Koch, M.D., et al., Pseudomonas aeruginosa distinguishes surfaces by stiffness using retraction of type IV pili. Proceedings of the National Academy of Sciences, 2022. 119(20): p. e2119434119.

9. Persat, A., et al., The mechanical world of bacteria. Cell, 2015. 161(5): p. 988–997.

10. Ribbe, J., et al., Role of Cyclic Di-GMP and Exopolysaccharide in Type IV Pilus Dynamics. J Bacteriol, 2017. 199(8).

11. Laventie, B.J., et al., A Surface-Induced Asymmetric Program Promotes Tissue Colonization by Pseudomonas aeruginosa. Cell Host Microbe, 2019. 25(1): p. 140–152.e6.

12. Yusuf, A.O., Z.K. Modi, and M.D. Koch, Rapid activation of dormant type IV pili enables a dispersal–infection tradeoff in environments with fluctuating nutrients. bioRxiv, 2026: p. 2026.04.23.720421.

13. Haque, F., et al., Pil-Chp orchestrates a multi-level regulatory system to control type IV pilus dynamics in Pseudomonas aeruginosa. bioRxiv, 2026: p. 2026.09.03.749161.

14. Schavemaker, P.E. and M. Lynch, Flagellar energy costs across the tree of life. eLife, 2022. 11: p. e77266.

15. Guo, S., et al., PilY1 regulates the dynamic architecture of the type IV pilus machine in Pseudomonas aeruginosa. Nature Communications, 2024. 15(1): p. 9382.

16. Porsch, E.A., et al., Pathogenic determinants of Kingella kingae disease. Frontiers in Pediatrics, 2022. Volume 10 - 2022.

17. Kus, J.V., et al., Significant differences in type IV pilin allele distribution among Pseudomonas aeruginosa isolates from cystic fibrosis (CF) versus non-CF patients. Microbiology, 2004. 150(5): p. 1315–1326.

18. Qiu, H. and W. Dai, Type IV PilD mutant stimulates the formation of persister cells in Pseudomonas aeruginosa. Journal of Antimicrobial Chemotherapy, 2025. 80(4): p. 1031–1036.

19. Bischof, L.F., et al., The Type IV Pilus Assembly ATPase PilB of Myxococcus xanthus Interacts with the Inner Membrane Platform Protein PilC and the Nucleotide-binding Protein PilM. J Biol Chem, 2016. 291(13): p. 6946–57.

20. Chang, Y.W., et al., Architecture of the type IVa pilus machine. Science, 2016. 351(6278): p. aad2001.

21. Sun, H., D.R. Zusman, and W. Shi, Type IV pilus of Myxococcus xanthus is a motility apparatus controlled by the frz chemosensory system. Curr Biol, 2000. 10(18): p. 1143–6.

22. Imhaus, A.F. and G. Duménil, The number of Neisseria meningitidis type IV pili determines host cell interaction. Embo j, 2014. 33(16): p. 1767–83.

23. Kraus-Römer, S., et al., External Stresses Affect Gonococcal Type 4 Pilus Dynamics. Frontiers in Microbiology, 2022. Volume 13 - 2022.

24. Eriksson, J., et al., Characterization of motility and piliation in pathogenic Neisseria. BMC Microbiol, 2015. 15: p. 92.

25. Holz, C., et al., Multiple pilus motors cooperate for persistent bacterial movement in two dimensions. Phys Rev Lett, 2010. 104(17): p. 178104.

26. Ellison, C.K., et al., Retraction of DNA-bound type IV competence pili initiates DNA uptake during natural transformation in Vibrio cholerae. Nat Microbiol, 2018. 3(7): p. 773–780.

27. Zhang, W., et al., Crash landing of Vibrio cholerae by MSHA pili-assisted braking and anchoring in a viscoelastic environment. eLife, 2021. 10: p. e60655.

28. Ellison, C.K., et al., Obstruction of pilus retraction stimulates bacterial surface sensing. Science, 2017. 358(6362): p. 535–538.

29. Iarocci, J., et al., <i>In situ</i> architecture of the Tad pilus machine in <i>Caulobacter crescentus</i>. mBio, 2026. 17(5): p. e00111–26.

30. Ellison, C.K., et al., Acinetobacter baylyi regulates type IV pilus synthesis by employing two extension motors and a motor protein inhibitor. Nature Communications, 2021. 12(1): p. 3744.

31. Ellison, C.K., et al., Real-time microscopy and physical perturbation of bacterial pili using maleimide-conjugated molecules. Nat Protoc, 2019. 14(6): p. 1803–1819.

32. Lynch, M. and G.K. Marinov, The bioenergetic costs of a gene. Proceedings of the National Academy of Sciences, 2015. 112(51): p. 15690–15695.

33. Winsor, G.L., et al., Enhanced annotations and features for comparing thousands of Pseudomonas genomes in the Pseudomonas genome database. Nucleic Acids Res, 2016. 44(D1): p. D646–53.

34. Craig, L., et al., Type IV pilus structure by cryo-electron microscopy and crystallography: implications for pilus assembly and functions. Molecular cell, 2006. 23(5): p. 651–662.

35. Craig, L., M.E. Pique, and J.A. Tainer, Type IV pilus structure and bacterial pathogenicity. Nature Reviews Microbiology, 2004. 2(5): p. 363.

36. Koch, M.D., et al., Competitive binding of independent extension and retraction motors explains the quantitative dynamics of type IV pili. Proceedings of the National Academy of Sciences, 2021. 118(8).

37. McCallum, M., et al., The molecular mechanism of the type IVa pilus motors. Nature Communications, 2017. 8(1): p. 15091.

38. McCallum, M., et al., Multiple conformations facilitate PilT function in the type IV pilus. Nature Communications, 2019. 10(1): p. 5198.

39. Wu, H., et al., Torque-speed relationship of the flagellar motor with dual-stator systems in Pseudomonas aeruginosa. mBio, 2024. 15(12): p. e0074524.

40. Lai, Y.W., et al., Evolution of the Stator Elements of Rotary Prokaryote Motors. J Bacteriol, 2020. 202(3).

41. Talà, L., et al., Pseudomonas aeruginosa orchestrates twitching motility by sequential control of type IV pili movements. Nature Microbiology, 2019. 4(5): p. 774–780.

42. Ellison, C.K., et al., A bifunctional ATPase drives tad pilus extension and retraction. Science Advances, 2019. 5(12): p. eaay2591.

43. Kilmury, S.L.N., et al., Hyperpiliation, not loss of pilus retraction, reduces <i>Pseudomonas aeruginosa</i> pathogenicity. Microbiology Spectrum, 2025. 13(4): p. e02558–24.

44. Yang, L., et al., In situ growth rates and biofilm development of Pseudomonas aeruginosa populations in chronic lung infections. J Bacteriol, 2008. 190(8): p. 2767–76.

45. Xia, X., Evolution of Translational Machinery in Fast- and Slow-Growing Bacteria. Microorganisms, 2026. 14(2): p. 377.

46. Li, S., et al., A quantitative study of the division cycle of Caulobacter crescentus stalked cells. PLoS Comput Biol, 2008. 4(1): p. e9.

47. Li, S., et al., Temporal Controls of the Asymmetric Cell Division Cycle in Caulobacter crescentus. PLOS Computational Biology, 2009. 5(8): p. e1000463.

48. Whitworth, D.E., et al., Phosphate Acquisition Components of the <i>Myxococcus xanthus</i> Pho Regulon Are Regulated by both Phosphate Availability and Development. Journal of Bacteriology, 2008. 190(6): p. 1997–2003.

49. Treuner-Lange, A., et al., Tight-packing of large pilin subunits provides distinct structural and mechanical properties for the Myxococcus xanthus type IVa pilus. Proc Natl Acad Sci U S A, 2024. 121(17): p. e2321989121.

50. Fronzes, R., H. Remaut, and G. Waksman, Architectures and biogenesis of non-flagellar protein appendages in Gram-negative bacteria. Embo j, 2008. 27(17): p. 2271–80.

51. Skerker, J.M. and L. Shapiro, Identification and cell cycle control of a novel pilus system in Caulobacter crescentus. Embo j, 2000. 19(13): p. 3223–34.

52. Sonani, R.R., et al., Tad and toxin-coregulated pilus structures reveal unexpected diversity in bacterial type IV pili. Proc Natl Acad Sci U S A, 2023. 120(49): p. e2316668120.

53. Cooley, B.J., et al., The extracellular polysaccharide Pel makes the attachment of P. aeruginosa to surfaces symmetric and short-ranged. Soft Matter, 2013. 9(14): p. 3871–3876.

54. Fernandez, N.L., et al., Vibrio cholerae adapts to sessile and motile lifestyles by cyclic di-GMP regulation of cell shape. Proc Natl Acad Sci U S A, 2020. 117(46): p. 29046–29054.

55. Kaiser, D., M. Robinson, and L. Kroos, Myxobacteria, polarity, and multicellular morphogenesis. Cold Spring Harb Perspect Biol, 2010. 2(8): p. a000380.

56. Drescher, K., et al., Architectural transitions in <i>Vibrio cholerae</i> biofilms at single-cell resolution. Proceedings of the National Academy of Sciences, 2016. 113(14): p. E2066–E2072.

57. Harris, L.K. and J.A. Theriot, Relative Rates of Surface and Volume Synthesis Set Bacterial Cell Size. Cell, 2016. 165(6): p. 1479–1492.

58. Conrad, Jacinta C., et al., Flagella and Pili-Mediated Near-Surface Single-Cell Motility Mechanisms in P. aeruginosa. Biophysical Journal, 2011. 100(7): p. 1608–1616.

59. Jin, F., et al., Bacteria use type-IV pili to slingshot on surfaces. Proceedings of the National Academy of Sciences, 2011. 108(31): p. 12617–12622.

60. Simsek, A.N., et al., Bacteria Tune a Trade-off between Adhesion and Migration to Colonize Surfaces under Flow. PRX Life, 2024. 2(2): p. 023003.

61. Persat, A., et al., Type IV pili mechanochemically regulate virulence factors in Pseudomonas aeruginosa. Proc Natl Acad Sci U S A, 2015. 112(24): p. 7563–8.

62. Wolfgang, M.C., et al., Coordinate regulation of bacterial virulence genes by a novel adenylate cyclase-dependent signaling pathway. Dev Cell, 2003. 4(2): p. 253–63.

63. Siryaporn, A., et al., Surface attachment induces <i>Pseudomonas aeruginosa</i> virulence. Proceedings of the National Academy of Sciences, 2014. 111(47): p. 16860–16865.

64. Klausen, M., et al., Biofilm formation by Pseudomonas aeruginosa wild type, flagella and type IV pili mutants. Mol Microbiol, 2003. 48(6): p. 1511–24.

65. Koch, M.D., et al., Competitive binding of independent extension and retraction motors explains the quantitative dynamics of type IV pili. Proceedings of the National Academy of Sciences, 2021. 118(8): p. e2014926118.

66. Gold, V.A.M., et al., Structure of a type IV pilus machinery in the open and closed state. eLife, 2015. 4: p. e07380.

67. Bulyha, I., et al., Regulation of the type IV pili molecular machine by dynamic localization of two motor proteins. Mol Microbiol, 2009. 74(3): p. 691–706.

68. Kilmury, S.L.N. and L.L. Burrows, The Pseudomonas aeruginosa PilSR Two-Component System Regulates Both Twitching and Swimming Motilities. mBio, 2018. 9(4).

69. Kilmury, S.L. and L.L. Burrows, Type IV pilins regulate their own expression via direct intramembrane interactions with the sensor kinase PilS. Proc Natl Acad Sci U S A, 2016. 113(21): p. 6017–22.

70. Rijal, A. and P.D. Curtis, Type IV pilin regulation: a transcriptional overview. Crit Rev Microbiol, 2026. 52(1): p. 36–63.

71. Christman, N.D., et al., The stoichiometry of minor-to-major pilins regulates the dynamic activity of the type IVa competence pilus in Vibrio cholerae. PLoS Genet, 2026. 22(6): p. e1012188.

72. Modi, Z.K., et al., Type IV pilus length determines virulence by regulating a putative subpopulation of non-contributing filaments in Pseudomonas aeruginosa. Communications Biology, 2026.

