## Supplementary Data for "The construction and operation of type IV pili impose a variable energetic burden across phylogenetically distant bacteria"

[#] Equal contribution

**Supplementary Data**

To quantify T4P machine number and PilA monomer density in individual cells, we used fluorescent labeling of the major pilin PilA using a *pilA*-A86C point mutant and Cysteine-maleimide click chemistry with an Alexa488-maleimdie dye to visualize pili [1, 2].

**Estimation of total number of pilus machines per cell.**

To estimate the total number of T4P machines, we used a retraction deficient strain (deletion of both retraction motors, *pilTU*) to count the number of pilus fibers per cell. Since pilus fibers do not retract in this mutant, T4P machines can be identified by the presence of a pilus (Supplementary Figure 1A). Among a total of *N* = 126 cells, we found that ~ 98% of cells do have at least one pilus, a typical cell has three pili, and the maximum of pili per cell was nine.

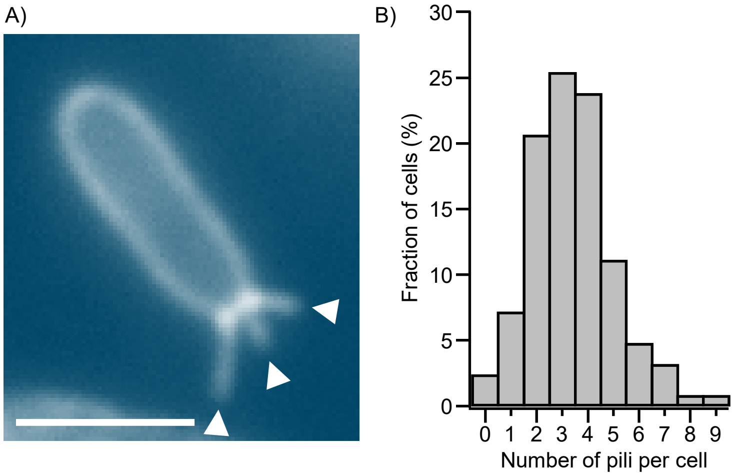


**Supplementary Figure 1. Estimating the number of T4P machines in single cells. A)** Image of a retraction deficient *pilTU* mutant with fluorescently labeled pili. Individual pili are marked with white triangles. Scale bar: 2 μm. **B)** Histogram of the number of pili per cell from N = 126 cells of three biological replicates.

**Estimation of the total number of PilA monomers per cell.**

Using videos of pilus extension and retraction, we previously demonstrated that the local concentration of pilin monomers at the based of a dynamic pilus machine changes during plus extension and retraction (Supplementary Figure 2A,B) [3]. Instead of measuring just the local intensity change at the base of the pilus, here we measured the total cell body fluorescence, representing all labeled PilA monomers. Comparing the change of total cell body fluorescence to the length of the extended pilus shows that both are inversely correlated, as expected (Supplementary Figure 2C). We further show that there is no hysteresis between extension and retraction, demonstrating that brightness loss/gain in the cell directly correlates with pilus length (Supplementary Figure 2D). Next, we analyzed the total change of cell body brightness for the maximum extension of N = 27 pili of different length from 15 cells total (Supplementary Figure 2E). A linear fit demonstrates a clear correlation of cell body brightness change to pilus length across cells. Since the absolute number of PilA monomers likely differs for cells with different length (assuming that cells keep the total inner membrane concentration of PilA per unit area constant), we normalized the brightness change by cell length. The density of PilA in the assembled pilus fiber is approximately one PilA per nanometer [4, 5]. Using this value, we estimated the number of PilA monomers in each pilus fiber, and from this the total number of PilA monomers that is equivalent to the total cell brightness. Adjusting for optical effects (see below), we estimate that an average cell has ~13,000 PilA monomers per micrometer of cell length, i.e., a typical L = 3 μm long cell has ~ 39,000 PilA total monomers.

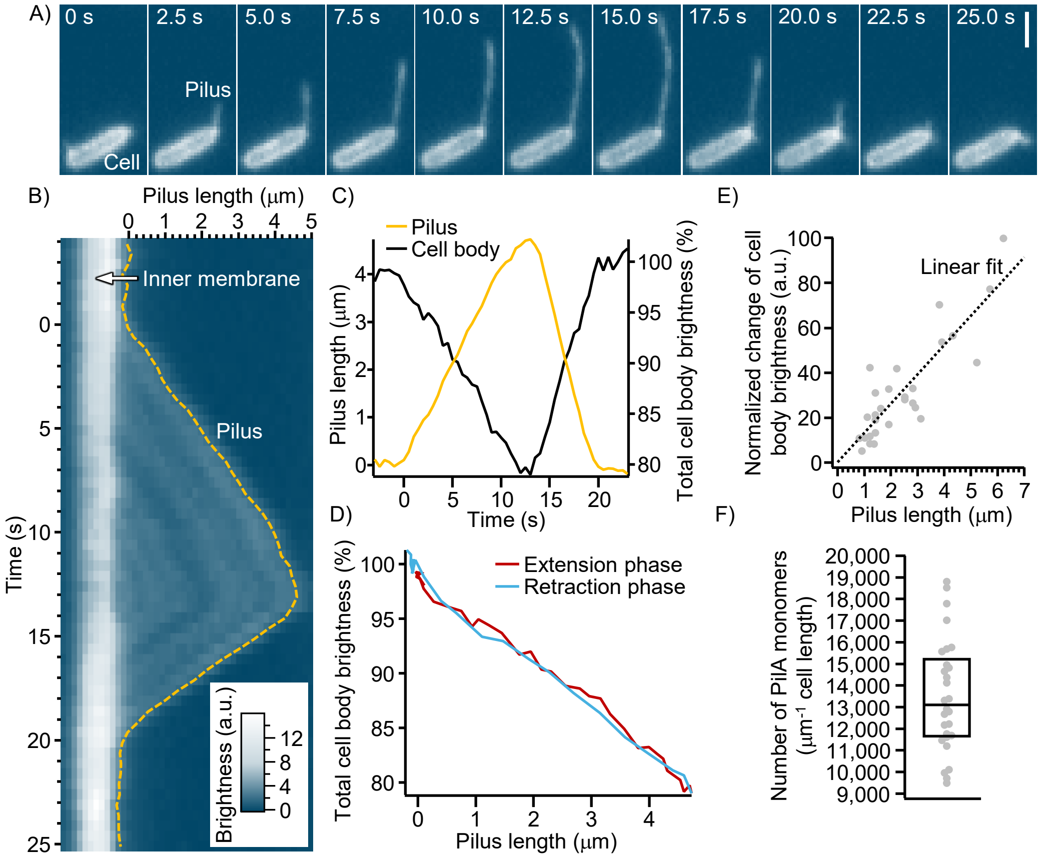


**Supplementary Figure 2. Estimating the number of PilA monomers per cell. A)** Fluorescence image time series of a single long pilus extending from the pole of the cell. Scale bar: 2 μm. **B)** Kymograph of the fluorescence intensity of a fixed line drawn along the length of the pilus showing pilus extension and retraction and a resulting change of PilA monomer concentration in the inner membrane at the base of the pilus. **C)** Correlation of changes in the total cell body fluorescence (labeled PilA monomer in the inner membrane) vs the current length of the extended pilus. **D)** The change of cell body brightness as a function of the maximum length of the pilus during the extension and retraction phase. **E)** Analysis of the total change of cell body brightness as a function of pilus lengths for N = 27 pili from 15 cells. The change of cell body brightness was normalized to the cell length. A linear fit was added to confirm a linear relationship between pilus extension length and changes in the total PilA concentration in the inner membrane (cell body fluorescence). **F)** Estimate for the number of PilA monomers in one cell, normalized to cell length.

### Supplementary Methods

#### PilA abundance correction factor to compensate for optical sectioning

Imaging of pilus fibers requires a high magnification, high numerical aperture microscope lens. Such lenses capture a limited optical depth which results in sectioning as shown in Supplementary Figure 1A and Supplementary Figure 2A. To estimate the total amount of PilA monomer in the cell, we correlated changes of the total cell body brightness with the length of extended pili. Since the images are optically sectioned, this underestimated the total protein count. Since the optical parameters of our microscope are well-known, we here calculate a correction factor that allows us to estimate the total monomer count from the optical sectioned images.

The cell was approximated as a cylinder of radius $r = 0.3$μm and cylindrical length $L = 2.4$μm with two hemispherical caps (total length $3$ μm). The captured fractions of fluorescence light of the cylindrical and spherical end cap parts captured by the microscope can be calculated separately using the following integral equations:

$$f_{\text{cyl}} = \frac{1}{\pi}\int_0^{\pi} \exp\left(\frac{(r\sin(\phi))^2}{2\sigma_z^2}\right) d\phi \approx 0.69,$$

$$f_{\text{cap}} = \frac{1}{2}\int_0^{\pi} \exp\left(\frac{(r\cos(\theta))^2}{2\sigma_z^2}\right) \sin(\theta)\, d\theta \approx 0.40$$

$\sigma_z = 211$nm corresponds to the full width half maximum (FWHM) of the gaussian point spread function (PSF) defined by $\text{FWHM}_z \approx \frac{2\lambda}{\text{NA}^2}$ = 497 nm. Here, the numerical aperture of our objective lens is NA = 1.45 and the emission maximum of the Alexa 488 dye is $\lambda = 519nm$.

Weighting of both factors by their relative surface area fractions (80% of the surface is the cylinder and 20% of the surface are the end caps) gives the overall effective capture factor

$$f_{\text{total}} = 0.8 \times 0.69 + 0.2 \times 0.40 = 0.63.$$

The total PilA protein count in the whole cell was obtained by dividing the measured and normalized cell body brightness change to pilus length ratio by $f_{\text{total}} = 0.63$ (see above).

### Supplementary References


1. Koch, M.D., et al., *Competitive binding of independent extension and retraction motors explains the quantitative dynamics of type IV pili.* Proceedings of the National Academy of Sciences, 2021. **118**(8).
2. Ellison, C.K., et al., *Real-time microscopy and physical perturbation of bacterial pili using maleimide-conjugated molecules.* Nature protocols, 2019. **14**(6): p. 1803-1819.
3. Koch, M.D., et al., *Pseudomonas aeruginosa distinguishes surfaces by stiffness using retraction of type IV pili.* Proceedings of the National Academy of Sciences, 2022. **119**(20): p. e2119434119.
4. Craig, L., M.E. Pique, and J.A. Tainer, *Type IV pilus structure and bacterial pathogenicity.* Nature Reviews Microbiology, 2004. **2**(5): p. 363.
5. Craig, L., et al., *Type IV pilus structure by cryo-electron microscopy and crystallography: implications for pilus assembly and functions.* Molecular cell, 2006. **23**(5): p. 651-662.
